# Haplotype-based eQTL mapping finds evidence for complex gene regulatory regions poorly tagged by marginal SNPs

**DOI:** 10.1101/314229

**Authors:** Robert Brown, Sriram Sankararaman, Bogdan Pasaniuc

## Abstract

**Motivation:** Expression quantitative trait loci (eQTLs), variations in the genome that impact gene expression, are identified through eQTL studies that test for a relationship between single nucleotide polymorphisms (SNPs) and gene expression levels. These studies typically assume an underlying additive model. Non-additive tests have been proposed, but are limited due to the increase in the multiple testing burden and are potentially biased by filtering criteria that relies on marginal association data. Here we propose using combinations of short haplotypes instead of SNPs as predictors for gene expression. Essentially, this method looks for genomic regions where haplotypes have different effect sizes. The differences in effect can be due to multiple genetic architectures such as a single SNP, a burden of rare SNPs, multiple SNPs with independent effect or multiple SNPs with an interaction effect occurring on the same haplotype.

**Results:** Simulations show that when haplotypes, rather than SNPs, are assigned non-zero effect sizes, our method has increased power compared to the marginal SNP method. In the GEUVADIS gene expression data, our method finds 101 more eGenes than the marginal method (5,202 vs. 5,101). The methods do not have full overlap in the eGenes that they find. Of the 5,202 eGenes found by our method, 707 are not found by the marginal method—even though it has a lower significance threshold. This indicates that many genes have regulatory architectures that are not well tagged by marginal SNPs and demonstrates the need to better model alternative archi-tectures.

## 1. Introduction

Expression quantitative trait loci (eQTLs) are genetic variants that regulate gene expression. Association scans to find eQTLs have been successfully applied to multiple datasets to find many eGenes, genes with at least one eQTL regulating their expression. These studies have shown that much of the variation in gene expression is heritable [1, 2, 3, 4] and that the genetic architectures that regulate expression are often found near the gene they regulate [5, 6]. The GEUVADIS [7] and GTEx [8] projects have publically provided genotype and expression data for hundreds of individuals across multiple tissues. This data has been successfully used to probe how genetic variation influences complex diseases through gene expression regulation [9, 10, 11, 12].

The standard test of association assumes an underlying additive model [13, 14] that relates genotype to expression level using a marginal single nucleotide polymorphism (SNP) test. eQTLs found through studies using this model may have independent additive effects, but could also have more complicated interactions that are only tagged by the marginal SNP test. The marginal SNP test does not account for the presence of multiple eQTLs existing for the same eGene [15] or for interactions between eQTLs for the same gene [16]. Recent work [17, 18, 19, 20] has searched for evidence of eQTLs arising from SNPxSNP interactions where one SNP moderates the effect of another. Such interactions are known to exist in yeast [21], but their relevance to gene expression in humans has been difficult to ascertain [22, 23, 24]. Other work has found evidence of haplotype effects when using a likelihood model for short haplotypes comprised of 10 SNPs [25] and for haplotypes defined by regions with a high likelihood of affecting gene expression [26]. However, these methods work for regulatory regions and must leverage functional predictions in order to reduce the computational burden to a small fraction of the genome. This makes their extension to complex phenotype studies problematic since they are restricted to only testing haplotypes within a megabase of genes.

In this work we present a new method (HapSet) for investigating haplotype effects on phenotype. While our method can be applied genome-wide with only a small increase in the computational burden, we apply it to gene expression data in order to test it against thousands of phenotypes. We hypothesize that short haplotypes each have their own specific independent effect on gene expression. The HapSet method divides haplotypes from a 10 kb region into all possible two set combinations and looks for a difference in the average effect size between the sets. A short region is selected so that the population can be charactorized by a small number of haplotypes. This test allows each haplotype to have its own genetic regulatory architecture and effect size and only assumes that there is no interaction between haplotypes. In order to use the marginal SNP method as a subset of our method, we force our method to include haplotype sets defined by the presence or absence of an alternate allele at specific SNP locations. Our method is not biased by filtering or variant selection that utilize marginal test statistics since it determines haplotype sets independent of the expression data. We compute significance thresholds for the haplotype set tests in order to control the family-wise error rate (FWER) at desired levels.

We simulate gene expression assuming five underlying architectures: single causal SNPs, multiple causal SNPs, SNPxSNP interactions, haplotypes with non-zero effects and a null model. Our simulations show that both methods maintain a 0.05 FWER under the null model. With the common SNP-based simulated architectures, the marginal SNP method has slightly higher power due to a less stringent significance threshold. We expect this result since the SNP method is a subset of the HapSet method. For example, when a single common SNP accounts for 5% of the variance in phenotype, the SNP method has 78% power while the HapSet method has 71%. However, when the underlying model is based on a random set of haplotypes assigned the same non-zero effect size, the HapSet method has 71% power compared to the SNP method that only has 56%. This demonstrates that there is insufficient SNP density to tag many of the possible haplotype combinations. We apply our method to find eGenes with the GEUVADIS data using expression and genotype data from 373 Europeans individuals and find evidence for many genes regulated by complex genetic architectures. Of the 18,621 genes in the data, the HapSet method finds 707 eGenes where the marginal SNP method had no significant association despite having a lower significant threshold. This demonstrates that for these 707 genes, there is likely to be a complex regulatory architecture that is poorly tagged by marginal SNPs. In these genes, the top HapSet association on average explains 6% of the variance in gene expression while the top (non-significant) marginal SNP only explains 3% of the variance. The marginal SNP method finds 606 eGenes not identified by the HapSet method. Since the marginal SNP method is a subset of the HapSet method, the HapSet method did not identify these eGenes due to its more stringent significance threshold. However, the top associations both explained 5% of the variance in expression. In addition to the eGenes found by only one of the methods, there are 4,495 eGenes where both HapSet and the marginal SNP method find significant associations with the top associations on average each explaining 10% of the variance in expression.

## 2. Methods

A SNP represents variation at one specific location in the genome. Haplotypes represent variation across a set of successive SNPs on a single chromosome. Generally, a small number of haplotypes are representative of the majority of the haplotypes in a sample when looking at a small region. Each of these haplotypes can have different effect sizes depending on the genetic architectures occurring on the haplotypes. Our proposed method seeks to maximize the difference between the frequency-weighted mean effect size on gene expression of haplotypes within a defined set (a HapSet) and those not in the set. Once a set is defined, haplotypes from individuals are identified as either in or out of the set, and a pseudo-genotype, defined as the number of haplotypes an individual carries that are within the set, is assigned. Gene expression can then be regressed on this pseudo-genotype to estimate the mean difference in effect size of haplotypes in and out of the set.

### 2.1. Haplotype effect model

We divide the genome up into *S* equal length sections. We assume that the *i^th^* haplotype in the *s^th^* section has its own specific and independent effect *β_si_* on gene expression *y*

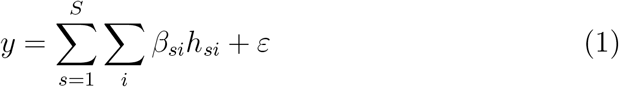

Here *y* is the gene expression for an individual, *h_si_* indicates the number of the *i^th^* haplotype in the *s^th^* section that an individual carries (either 0, 1 or 2) and 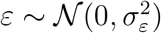. When only interested in finding associations between *y* and any of the haplotypes in a region *s*, we can rewrite our model (Equation 2). Here *ε* now includes the variance due to all of the haplotypes not in region *s* as well as environmental noise.

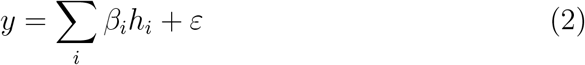

### 2.2. Marginal SNP method

The marginal SNP method for identifying an association between a SNP and expression fits the following additive model

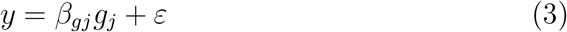

where *g_j_* is the number of alternate alleles an individual carries at the *j^th^* SNP. Assuming the model in Equation 2, *β_g_j__* will be the difference between the frequency weighted average effect of haplotypes containing an alternate allele at SNP *j* and the average effect of haplotypes that carry the reference allele:

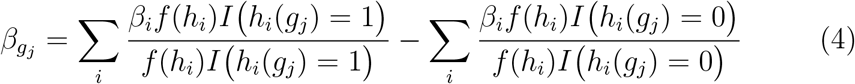

Here *I* (*h_i_*(*g_j_*) = 1 is equal to 1 if *h_i_* contains an alternate allele at the *j^th^* SNP (*g_j_*) otherwise it is 0. Similarly I (*h_i_*(*g_j_*) = 0) is equal to 1 if *h_i_* contains a reference allele at the *j^th^* SNP (*g_j_*) otherwise it is 0. The frequency of haplotype *h_i_* is given by *f* (*h_i_*).

### 2.3. Haplotype set method

The haplotype set (HapSet) method fits an additive model similar to Equation 3

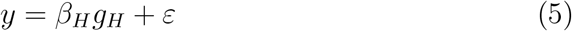

where *g_H_* is a pseudo-genotype for the number of haplotypes an individual carries that are members of a set *H* of haplotypes (see Section 2.5). *β_H_* is the difference between the average effect size of haplotypes in set *H* and the average effect size of haplotypes not in set *H* (i.e. in set *H^c^*)

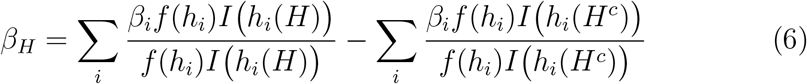

Here *h_i_*(*H*) and *h_i_*(*H^c^*) are equal to 1 if haplotype *h_i_* is a member of set *H* (or *H^c^*) and 0 otherwise.

### 2.4. Testing for significant associations

A standard measure of association is the Wald statistic for expression levels *y* of *n* individuals and genotypes (or pseudo-genotypes) *g* with 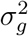 variance

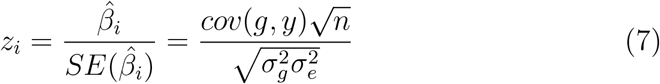

which asymptotically follows a normal distribution with variance 1 and a non-centrality parameter given by

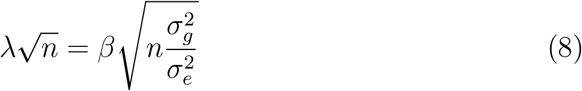

Since the 𝔼(*βȃ*) = *β* under the marginal SNP method, testing the SNP with the largest *β_g_* after standardizing *g_j_* will result in the largest association statistic for the marginal SNP test. Likewise, for the HapSet method, a set *H* that maximizes *β_H_* for a standardized *g_H_* will maximize the association statistic for the HapSet method.

### 2.5. Determining haplotype sets

Since there is no way to know *a priori* the *H* that maximizes the difference, we test all possible *H*s from all 10 kb windows within 500 kb of a gene’s transcript start site. To determine the *H*s, we remove all SNPs with minor allele frequency *<* 0.05 and find all the haplotypes in a 10 kb region. We choose a small region so that a small number of haplotypes will be represenative of the majority of the haplotypes in the sample. We treat all haplotypes with frequency *<* 0.05 as a single haplotype. If there are more than 8 haplotypes in the group, we use the 7 most frequently occurring haplotypes and treat all the remaining haplotypes as the 8*^th^*haplotype. We then compute all possible ways to form two sets (*H* and *H^c^*). Using the phased data provided, for each *H* we encode for each individual the number of haplotypes they carry that are contained in *H* as *g_H_*. We compute a *g_H_* for each *H* of every 10 Kb region in the genome.

We force the marginal SNP method to be a subset of the HapSet method. For each SNP with minor allele frequency *>* 0.05, we define all haplotypes with an alternate allele at the SNP location to be in set *H* and all haplotypes with the reference allele at the SNP location to be in set *H^c^*.

### 2.6. Power simulations

We simulate four causal architectures using genotype data from chromosome 22 of the 373 Europeans in the GEUVADIS project. We use the non-transformed genotype data. We simulate using the following models where the proportion of the variance due to the underlying architecture is *h*^2^: null, single causal SNP, two causal SNPs, SNPxSNP interaction and a haplotype set model (Equations 9, 10, 11, 12 and 13 respectively). Causal SNPs (*g_C_*_1_ and *g_C_*_2)_ are randomly chosen from SNPs within 500 Kb of a transcription start site. All SNPs have minor allele frequency *>* 0.05 except when using a rare single causal SNP with frequency between 0.01 and 0.05. We fix all SNP or HapSet effect sizes to be 1 (*β_C_*_1_ = *β_C_*_2_ = *β_H_* = 1) and add noise to achieve the desired *h*^2^. The haplotype set model assumes that a random set of haplotypes *H* all have the same non-zero effect size. We draw the causal *H* from one of the *H*s determined in section 2.5 that is within 500 kb of a transcription start site. When analyzing how the methods perform when the simulated causal haplotypes are not present in the data, (masked causal haplotypes), we do not test any SNPs or *H*s from the 10kb region that contains the causal haplotypes. We then regress the simulated expression on all genotypes and *g_H_* s within 500 kb of the transcription start site. We simulate each trait 65,000 times for each *h*^2^ and the null model 650,000 times.

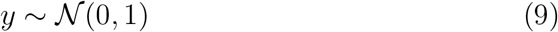

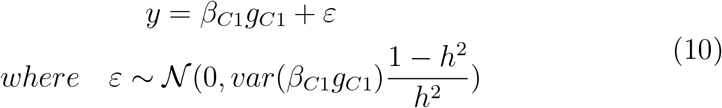

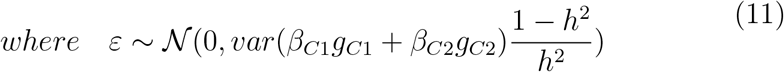

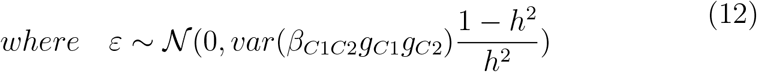

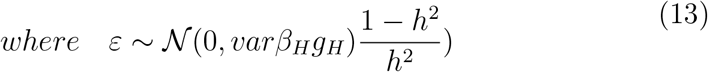

## 3. Results

### 3.1. Data for simulations and analysis

Our analyses are based on publically available genotype and lymphoblas-toid expression data of 373 European individuals provided by the GEUVADIS [7] project. Following the GEUVADIS project, we only use expression from genes that had *>* 0 quantifications in *>* 90% of individuals. We center and standardize the RPKM and PEER normalized gene expression levels.

### 3.2. Controlling the family-wise error rate

Using all of the *g_H_* values calculated from the chromosome 22 genotypes of the 373 European individuals in the GEUVADIS data and the genotype data for SNPs with minor allele frequency above 0.05, we compute significance thresholds using SLIDE [27]. Since eQTL testing only looks at predictors within 500 kb of each gene, we ensure a family-wise error rate (FWER) level for each gene by scaling the threshold for a 1 Mb region. For a 0.05 desired FWER, this results in a 2.0 × 10^−5^ significance threshold for our HapSet method. Running the same procedure using the genotype data for the SNP method, we estimate a per megabase significance threshold of 5.8×10^−5^. The results from the different FWER levels (see Table 1) suggest that the HapSet method needs to correct for approximately 3 times as many independent tests in comparison to the marginal SNP method.

**Table 1:** Per megabase significance thresholds estimated with SLIDE. For the 0.01, 0.05 and 0.1 FDRs analyzed, the HapSet method performs approximately 3 times as many independent tests as the SNP method.

|  | Marginal SNP Method | HapSet Method |
| --- | --- | --- |
| 0.01 FWER | $9.8 \times 10^{-6}$ | $3.3 \times 10^{-6}$ |
| 0.05 FWER | $5.8 \times 10^{-5}$ | $2.0 \times 10^{-5}$ |
| 0.10 FWER | $1.2 \times 10^{-4}$ | $4.0 \times 10^{-5}$ |

### 3.3. Power analysis

Our simulations show that the marginal SNP method and HapSet method are both well calibrated under the null with 0.043 and 0.046 discovery rates, respectively, when controlling for a 0.05 FDR. The power of both methods increases as the variance due to the underlying genetic architecture increases.

The discovery rate of the SNP method is slightly superior to that of the HapSet method when the underlying model is based on common SNPs (see Figure 1 and Table 2). However, if the underlying model is a single “rare” SNP with minor allele frequency between 0.01 and 0.05, the HapSet method outperforms the marginal SNP method. When the underlying model is a random set of haplotypes *H*, the HapSet method shows a substantial increase in power over the marginal SNP method (see Figure 2 and Table 2). For example, for an *h*^2^ = 0.05 and a haplotype set architecture, the SNP method has 56% power compared to 71% for the HapSet method (see Table 2). When the set of causal haplotypes is masked, the HapSet method still outperforms the marginal SNP method.

**Figure 1.**
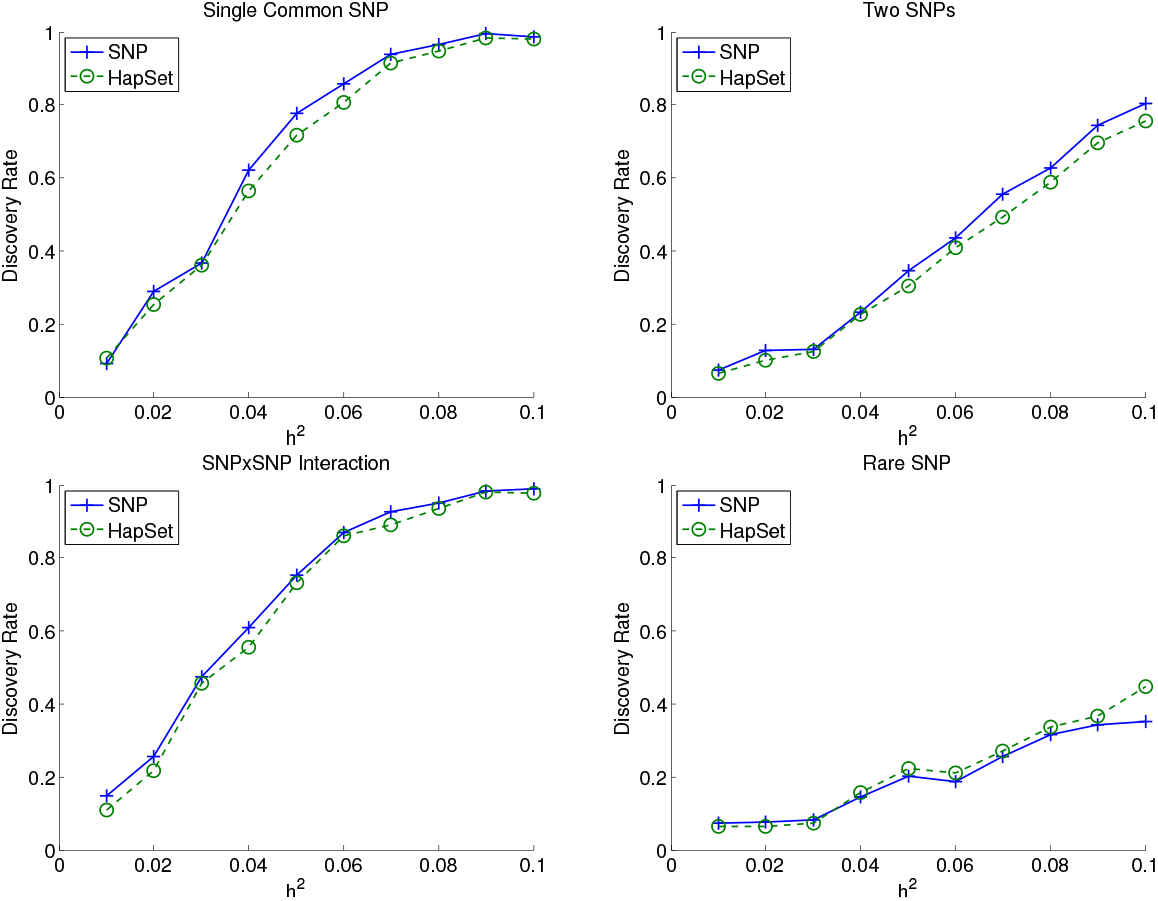
Discovery rate of the marginal SNP and HapSet methods with SNP-based simulated architectures. The proportion of variance due to the underlying genetic architecture is given by *h*^2^. The HapSet method slightly outperforms the SNP method for causal SNPs with minor allele frequencies between 0.01 and 0.05. Since the HapSet method has a more stringent significance threshold to control the FWER at 0.05, this indicates that the HapSet method is better tagging the causal SNP than the SNP method. For all other SNP-based causal architectures, the HapSet method performs slightly below the SNP method. Since the SNP method is a subset of the HapSet method, this is due only to the difference in significance thresholds.

**Figure 2.**
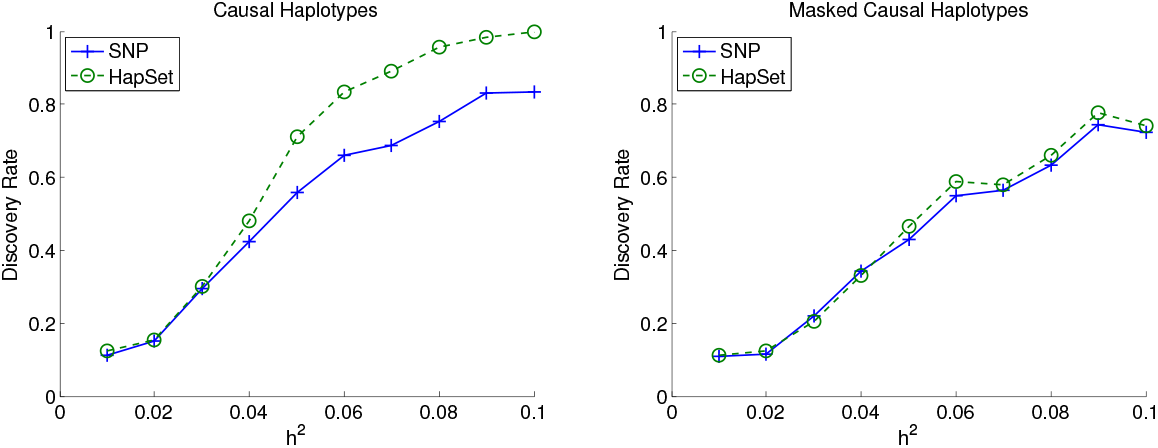
Discovery rate of the marginal SNP and HapSet methods with haplotype-based simulated architectures. When a random set of haplotypes has a non-zero effect size, the HapSet method has a large power advantage over the marginal SNP method. When the set of causal haplotypes is masked, the HapSet method slightly outperforms the SNP method.

**Table 2:**
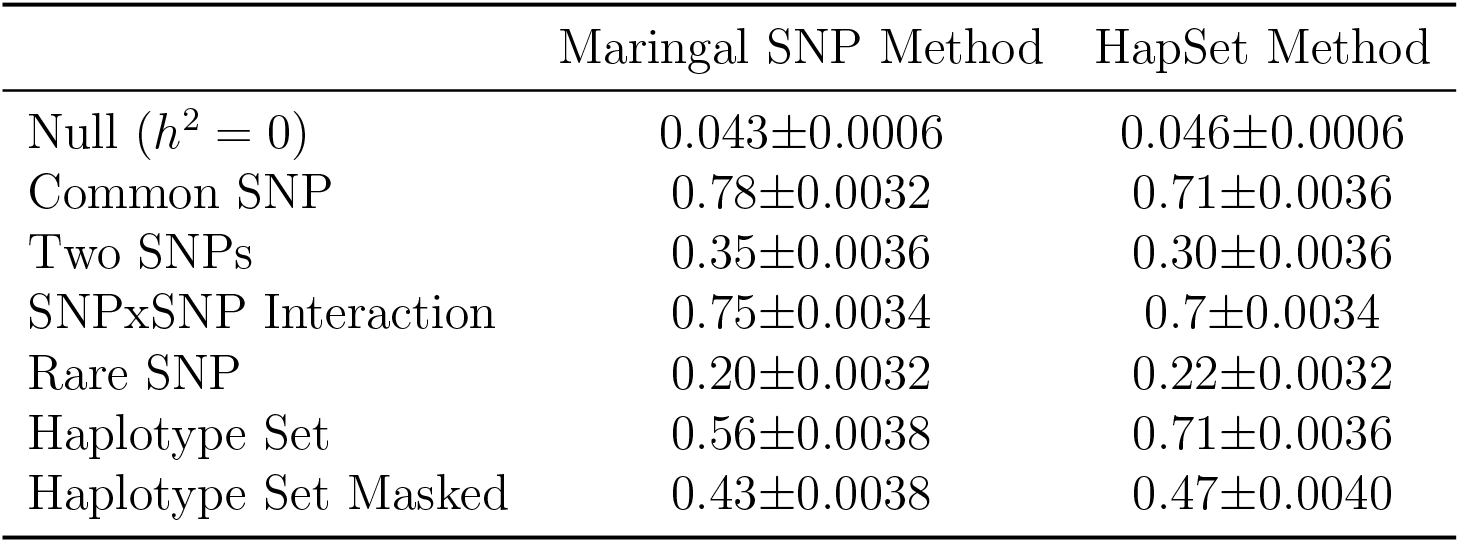
Discovery rate (±2*SE*) of the marginal SNP and HapSet methods when *h*^2^ = 0.05. The marginal SNP method outperforms the HapSet method when the underlying genetic architecture is based on SNPs. The exception is when the underlying architecture is based on a ‘rare’ SNP with allele frequency between 0.01 and 0.05. When the architecture is based on haplotype sets, the HapSet method strongly outperforms the SNP method. When the haplotype sets and SNPs within 10kb of the simulated casual haplotype set are not included in the data for testing (masked), the HapSet method still outperforms the SNP method. This indicates that there is not enough SNP density to tag some haplotype combinations.

### 3.4. Analysis of GEUVADIS data

We apply the two methods to the GEUVADIS gene expression data. Following the GEUVADIS analysis, we regress expression on the genotypes or HapSets (both standardized to mean 0 and variance 1) while controlling for the top three genotype-based PCs [7]. We use the 18,621 genes that passed the filtering criteria. We report in Table 3 the number of eGenes found by the SNP method and the HapSet method while controlling for different FWER levels.

**Table 3:**
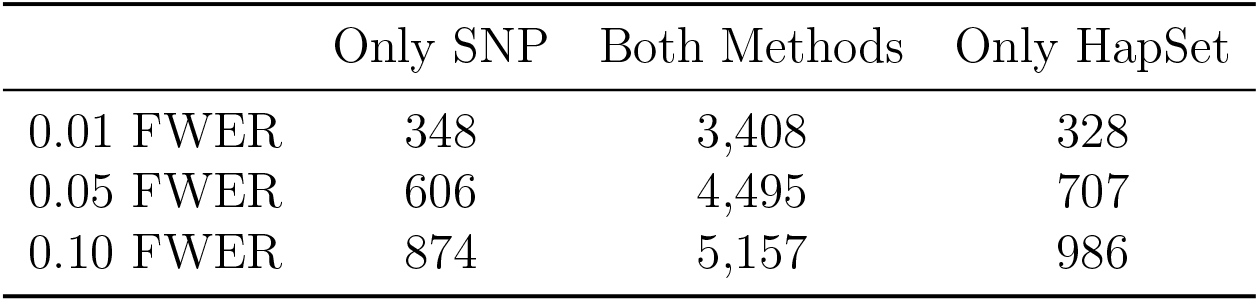
Number of eGenes identified by the marginal SNP and HapSet methods while controlling for a given FWER. The marginal SNP method identifies 20 more eGenes than the HapSet method when using a 0.01 FWER, but with the 0.05 and 0.1 FWERs, the HapSet method identifies 101 and 112 more eGenes respectively.

When controlling for a 0.05 FWER, the two methods found 4,495 overlapping eGenes (see Table 3). The HapSet method also found 707 eGenes that were not detectable using genotype data. Since the SNP method is a subset of the HapSet method, yet has a less stringent significance threshold, this indicates that marginal SNPs poorly tag the regulatory architecture for these genes. Conversely, the 606 eGenes found only by the SNP method represent eGenes that had p-values smaller than 5.8 × 10^−5^ (the SNP method threshold for a 0.05 FWER) but larger than 2.0 × 10^−5^ (the HapSet threshold for a 0.05 FWER). In total, there are 101 more eGenes identified by the HapSet method than by the marginal SNP method. Using a more stringent 0.01 FWER shows that the marginal SNP test identifies 20 more eGenes than the HapSet method. However, by allowing for a looser FWER of 0.1, the HapSet method again outperforms the marginal SNP method and identifies 112 more eGenes.

We evaluate the average squared effect sizes for the top SNP or HapSet for each eGene when controlling for a 0.05 FWER. The 4,495 eGenes found by both methods had mean squared effect sizes of 0.097 and 0.099 for the SNP and HapSet method, respectively. The 606 eGenes only found by the SNP method had effects sizes of 0.045 (SNP) and 0.045 (HapSet). The SNP method is a subset of the HapSet method but has a less stringent significance threshold, which explains why these eGenes were not found by the HapSet method but have the same mean squared effect size. The mean squared effect sizes for the eGenes found only by the HapSet method have the largest difference of 0.034 (SNP) and 0.055 (HapSet). This difference may indicate that there are underlying causal architectures that the HapSet method is capable of effectively tagging but the marginal SNP method cannot identify. We regress out the effect of the top associated HapSet for the 707 eGenes found only by the HapSet method and compute the mean squared effect of the same top (though not significant) SNP associations with the residuals. We observe a mean squared effect of 0.013. This indicates that the top SNPs were largely picking up on signal better captured by the HapSet method.

## 4. Discussion

In this work we present a new framework for finding regions of the genome with complex regulatory architectures that are poorly captured by a marginal SNP-based regression method. We assume that the two haplotypes an individual carries have additive non-interacting effects, but make no assumptions about the form of the regulatory architecture on each haplotype. We implement our method and demonstrate that it is well calibrated under the null hypothesis. We also show that it has little loss of power due to an increase in the multiple testing burden for common SNP architectures, but it has increased power for SNPs with *<* 0.05 minor allele frequency and haplotype-based architectures.

When we apply our method to real gene expression data, we see increased power to detect eGenes as compared to the standard marginal SNP method. Most importantly, we observe a large number of eGenes that are detected by the HapSet method yet undetected by the SNP method. Since the SNP method is a subset of the HapSet method, but with a less stringent significance threshold, this indicates that in many genes there are complex regulatory effects that are poorly captured by marginal SNP association methods. Our method gives no indication of the specific causal architectures underlying each of the eGenes found with the HapSet method. Further analyses must be performed in order to identify if there are multiple independent causal SNPs, a SNPxSNP interaction or a more complicated haplotype based effect. It is possible that our method may be identifying rare SNPs or un-typed genetic variation. Confounding from such variables has been observed in tests associating SNPxSNP interactions with gene expression [22, 23, 28].

Future work can implement a version of our method for identification of associated loci in complex traits. Many complex traits have GWAS loci that overlap known eQTLs [29], and applying the HapSet method to complex trait data is a natural extension. The second direction is to apply the HapSet method to fine mapping. Modeling more than 2 or 3 causal SNPs in a region for fine mapping can be computationally challenging [30, 31, 32, 33, 34, 35, 36]. However, knowing which combinations of haplotypes are associated, and to what degree, may improve efficient selection of SNPs and interactions between SNPs for testing.

